# Tomato Omics Research: Mapping Bibliometric Footprints of Global Research Trends, Collaboration Networks, and Impact Trajectories

**DOI:** 10.1101/2025.04.17.649284

**Authors:** Ratna Prabha, Amrender Kumar, Renu, Dhananjaya Pratap Singh

**Affiliations:** Agricultural Knowledge Management Unit (AKMU), ICAR-Indian Agricultural Research Institute, New Delhi 110 012 INDIA; Indian Council of Agricultural Research, Krishi Bhavan, New Delhi 110 001 INDIA; ICAR-Indian Institute of Vegetable Research, Shahanshahpur, Varanasi 221 305, INDIA

**Keywords:** Web of Science, Bibliometric analysis, Text mining, Network analysis, Tomato-Omics big data, Bioinformatics

## Abstract

Tomato is a globally vital crop and model system for fleshy fruit development, with profound agricultural and economic significance. Omics-based tomato research has revolutionized understanding of fruit quality traits, disease resistance mechanisms, and stress tolerance pathways reflecting insights for breeding innovations to address food security challenges. This bibliometric analysis on tomato omics delivered crucial insights into the evolution and impact of tomato omics research over two decades. Examining 1,702 Scopus-indexed publications, we have shown a remarkable acceleration in research output after 2017, signifying the field’s growing significance. Our findings revealed that Biochemistry, Genetics and Molecular Biology dominated the landscape with 1,054 publications, establishing the molecular foundation for tomato research. It identified key global research hubs, with the United States (450 publications) leading, followed by China (310) and Italy (171), while mapping eight distinct International collaboration networks that drive innovation. Through citation analysis, we have pinpointed transformative contributions in microRNA research and genome sequencing, with the field’s most influential paper garnering 2,570 citations. Co-authorship network visualization exposed 25 research clusters among 8,103 authors, highlighting prolific contributors including Fernie A.R. (39 publications) and Lucini L. (26). The keyword co-occurrence mapping revealed critical research priorities centered on stress responses, fruit quality, and plant immunity, with “Frontiers in Plant Science” emerging as the field’s premier publication venue (92 articles). Our analysis provided a roadmap of tomato omics research progression, offering strategic value for prioritization, collaboration development, and funding allocation to address pressing challenges in global food security and crop improvement.

## Introduction

Tomato (*Solanum lycopersicum* L.) being one of the most economically important vegetable crops globally, are rich in phytochemicals and nutrients like lycopene, potassium, iron, folate, and vitamin C along with other antioxidants such as beta-carotene, flavonoids, and phenolic compounds (Bhowmik et al, 2012; Toor et al, 2005; Borguini and Ferraz, 2009). With their versatility, tomatoes can be consumed raw or cooked, maintaining their nutritional value, thus making them valuable additions to a healthy diet (Bhowmik et al, 2012; Collins et al, 2022). Over 80% of commercially grown tomatoes are processed into products like juice, soup, and ketchup (Viuda-Martos et al, 2014), contributing to their widespread dietary consumption. The antioxidant-rich content of tomatoes and their products is linked to various health benefits (Bhowmik et al, 2012; Rao and Agrawal, 1999; Kearney et al, 2005; Collins et al, 2022). With the rapid advancements in omics technologies like genomics, transcriptomics, proteomics, and metabolomics since the early 2000s, tomato research has witnessed a remarkable surge in scientific output (Tilesi et al, 2021; Bento-Da-Silva et al, 2023). Bibliometric analysis provides a quantitative approach to map and evaluate the research trends, productivity, and impact in a particular field over time (Pritchard, 1969; Broadus, 1987). In the last two and a half decades (2001-2023), omics research on tomato has seen an exponential growth across the world. The completion of the tomato genome sequence in 2012 was a major milestone, providing a background and igniting more focused genomics and integrated multi-omics studies (Tomato Genome Consortium, 2012). Transcriptomics has been extensively used to elucidate gene expression patterns related to various biological processes, stress responses, and trait development in tomato (Kulshrestha and Parihar, 2020). Proteomics and metabolomics have complemented genomics and transcriptomics, enabling a systems biology approach to understand the complex molecular networks underlying tomato biology (Tang et al, 2020; Naik et al, 2023; Singh et al, 2023a,b; Singh et al, 2024). We first discussed a comprehensive overview on the theme, narrating its significance and then moved to exploring global trends in this area through bibliometric data analysis.

Our analysis on the bibliometric data on tomato omics studies from 2001 to 2023 identified under-explored areas or research gaps in tomato omics research and suggested for the need of the informed policy making and funding decisions to prioritize research, allocate resources, and develop strategies for addressing enhanced global competitiveness on tomato research and development.

### Omics Research in Tomato: Background Overview

Tomato research has benefited significantly from various -omics technologies, providing valuable insights into fruit quality, nutritional value, and stress responses. These technologies have revolutionized our understanding of tomato biology and accelerated breeding programs for improved varieties. Genomic technologies have been pivotal in studying tomato diversity, taxonomy, and trait mapping (De Vos et al, 2011; Granell et al, 2007). The completion of the tomato genome sequencing project has provided a high-quality assembly, serving as a foundation for sequencing-based approaches and accelerating gene identification and functional assignment (Sato et al, 2013; Sato and Tabata, 2016). High-scale sequencing methods have offered insights into the genetic architecture of tomato germplasm resources and evolutionary history (Tripodi, 2022; Wang et al, 2024).

The development of new genetic and genomic tools has increased the efficiency of tomato breeding, addressing challenges in horticultural production (Gramazio et al, 2018). Molecular markers and cloned resistance genes have enabled high-throughput screening for multiresistant varieties (Barone et al, 2009). The CRISPR/Cas9 system has been applied for genome editing, particularly for improving fruit quality and nutritional value (Scarano and Santino, 2023; Wang et al, 2020). These studies have resulted in the development of several databases and web resources to support tomato genomics research. Among them are the *i.* Tomato Functional Genomics Database (TFGD) that provides tools for data mining and analysis of large-scale functional genomics datasets (Fei et al, 2011), *ii.* Tomato Genomic Resources Database (TGRD) which allows interactive browsing of tomato genes, microRNAs, and genetic maps (Suresh et al, 2014) and *iii.* Sol Genomics Network (SGN) which contains biological data for Solanaceae species, including tomatoes (Bombarely et al, 2011).

Transcriptomics has revealed dynamic gene expression patterns, aiding in identifying candidate genes for desirable traits and enabling targeted breeding and genetic engineering (Vishwanath et al, 2024). It has been used to understand relationships between important traits, metabolism, and the development of next-generation breeding strategies (Vishwanath et al, 2024; Kusano, and Fukushima, 2013). Various methods have been employed for transcriptomic analysis in tomatoes, including 1) Quantitative PCR (qPCR) (Duan et al, 2022; Cheng et al, 2017), 2) RNA-Seq (Matas et al, 2013), 3) Laser microdissection coupled with RNA-Seq (Shinozaki et al, 2018), 4) High-throughput Gene Sequencing Methods (Ronnig, 2013) and 5) Microarrays. Such studies have identified numerous stress-responsive genes in this plant, providing insights into genetic mechanisms underlying stress responses (Iovieno et al, 2016; Diouf et al, 2020; Gan et al, 2024). Key transcription factor families, such as WRKY, MYB, and NAC, have been identified as playing pivotal roles in regulating plant responses to abiotic and biotic stresses (Jena et al, 2022; Mo et al, 2021; Fan et al, 2020).

Proteomics has contributed significantly to tomato research, offering insights into various aspects of tomato biology and development. It has been particularly useful in studying fruit development and identifying proteins involved in various physiological pathways (Luo et al, 2020). Proteomics technologies, especially when combined with other approaches, have been powerful tools in identifying functional proteins in tomato fruit tissues (Luo et al, 2020). Proteomic studies have complemented functional genomics and transcriptomic research, providing a comprehensive understanding of tomato growth, development, and stress responses (Momo et al, 2023; Sant’Ana et al, 2018). Advanced methodologies such as phosphoproteomics and isobaric tags have enabled the study of metabolic dynamics and regulations in tomatoes (Sant’Ana et al, 2018). Challenges in proteomic research include establishing methodologies that result in the best quality and wide-range representation of total proteins, especially when dealing with limited amounts of tissue (Vilhena et al, 2015). To address these challenges, optimized protocols for shotgun proteomics have been developed, facilitating the detection of a higher number of proteins and demonstrating reproducibility and robustness (Kilambi et al, 2016).

Metabolomics has been instrumental in understanding the relationship between important traits and metabolism in tomatoes, contributing to the improvement of taste, quality, stress resistance, and health-beneficial metabolites (Kusano and Fukushima, 2013; De Vos et al, 2011). It has facilitated the identification of metabolites associated with fruit quality, nutritional value, and the impact of storage and processing on these traits (De Vos et al, 2011; Ara et al, 2017). Recent advancements in mass spectrometry technology have enabled comprehensive, high-throughput profiling of metabolites in tomato plants, allowing for the simultaneous detection of a large number of metabolites and investigation of metabolic regulation affecting fruit productivity and quality (Iijima and Aoki, 2009; Carrari et al, 2006). The development of common metabolite databases, such as the Metabolome Tomato Database (MoTo DB), has provided comprehensive knowledge of the metabolite composition of tomato fruit (Moco et al, 2006).

Metabolomics has been particularly useful in studying stress responses in tomatoes as is evident from the studies of Singh et al, (2023a.b, 2024). It has characterized the effect of combined stresses like salinity and heat on tomato plant metabolism (Lopez-Delacalle et al, 2021) and detected metabolomic fluctuations in response to drought stress in tomato fruits (Asakura et al, 2022). These studies have revealed modulation of primary and secondary metabolism pathways under stress conditions, providing actionable knowledge for sustainable agriculture and food production (Chele et al, 2021). Overall, the integration of genomics, transcriptomics, proteomics, and metabolomics has significantly advanced our understanding of tomato biology, stress responses, and fruit quality. These technologies continue to play crucial roles in developing improved tomato varieties with enhanced agronomic traits, stress tolerance, and nutritional value.

### Bibliometric Analysis

Omics research in tomato is fitting case for bibliometric analysis. Such an analysis could provide valuable insights into research trends, collaboration patterns, and the impact of this field over time. The suitability for bibliometric analysis and potential interpretations of the data can be speculated keeping in view the rapidly evolving field of omics technologies in tomato research that have advanced significantly in recent years, making it an ideal subject for tracking research trends over time. Because of the interdisciplinary nature, tomato omics research spans genomics, transcriptomics, proteomics, and metabolomics, involving researchers from various disciplines. This diversity makes it interesting to analyze collaboration patterns and knowledge flow between different fields. Since the crop is commercially viable worldwide, the economic importance of tomatoes is globally accepted, which likely translates to a substantial body of research literature suitable for quantitative analysis. Furthermore, technological advancements in the field has evolved rapidly, making it interesting to study how new technologies have influenced research output and focus. In a similar way, the potentiality for interpretations of bibliometric data has been realized when we look at the research trends via publication volume over time, which could reveal the growth of tomato omics and identify periods of accelerated research activity. Analyzing keywords and abstracts could show how research focus has shifted between different omics approaches (e.g., from genomics to metabolomics) or specific research questions over time. An important shift in methodological approaches by tracing the adoption of new technologies or experimental approaches in publications could also be highlighted. Collaboration patterns could indicate co-authorship networks to reveal key research groups, institutions, and countries leading tomato omics research while analyzing author affiliations that could show the extent of global collaboration in this field and its evolution. Overall, the study can suggest trends, growth and limitations in research impact including citation and journal analysis which could reveal preferred journals and periodical shifts in publication strategies. It can also suggest funding patterns while analyzing major funding sources and how they’ve influenced research directions, technology adoption to show how newer technologies like RNA-Seq, CRISPR are adopted in tomato research. The bibliometric analysis can also comprehensively analyse the context of omics research (e.g., stress response, fruit quality) could reveal the main areas where these technologies are being applied in tomato research. The analysis could also indicate emergence of big data approaches that could become a trend towards publications mentioning machine learning or artificial intelligence for analyzing large-scale omics data. The approach can bridge basic and applied research in which the data might reveal increasing collaboration between academic institutions and commercial breeding companies, indicating a push towards practical applications of omics research and can lead to identify geographical shifts by showing changes in the global distribution of tomato omics research, potentially revealing emerging research hubs. By conducting a comprehensive bibliometric analysis, researchers could gain valuable insights into the evolution, current state, and future directions of omics research in tomatoes. This could guide future research efforts, inform funding decisions, and highlight areas where more attention might be needed.

## Materials and Methodology

### Data Collection

Clarivate Analytics’ Web of Science (WoS) and Elsevier’s Scopus are the major databases and citation indexes for general scientific literature, encompassing journal articles, conference proceedings, and books across various domains. While WoS predominantly covers natural sciences and engineering, Scopus has a broader scope, with a relatively higher emphasis on social sciences (Mongeon & Paul-Hus, 2016; Kumpulainen & Seppänen, 2022). WoS was the pioneer international bibliographic database and for a long time, it was the cornerstone for journal selection, research evaluation, and bibliometric analyses. It held a monopoly for over 40 years until the advent of Scopus in 2004, marking a significant shift in the landscape (Li et al, 2018; Baas et al, 2020). Scopus quickly emerged as a formidable alternative, boasting broader coverage and comparable reliability to WoS (Zhu and Liu, 2020; Harzing and Alakangas, 2016; Pranckutė, 2021).

Numerous comparisons have affirmed Scopus’s wider scope and unique coverage, rendering it more suitable for research evaluation and daily tasks (Wouters et al, 2015; Pranckutė, 2021). Scopus’s user-friendly interface, comprehensive author and institutional profiles, and robust impact indicators further underscore its practicality and accessibility compared to WoS. Additionally, Scopus’s single-database subscription model and transparent access to author and source information, including metrics, contribute to its broader societal impact (Pranckutė, 2021). For the current study, SCOPUS database was used for the bibliometric analysis. In the process of data collection, the search field utility was accessed in which the field was restricted to ‘Title’ (TITLE), ‘Abstract’ (ABS) and ‘Keywords’ (KEY). The queries were generated using the words ‘Tomato’ and ‘solanum lycopersicum’ along with important terms from the field of –omics. The possible combinations were queried using ‘AND’ and ‘OR’ logical operators in this process. The initial exclusion criteria on language, year of publication and document type were applied to filter the dataset. Further, the relevancy was checked of filtered publications to obtain the final set of data. All publications of duration 2001-2023 were considered, Document Types were limited to Article, Review, Book chapter, Book, Letter and Short survey while Source type were restricted to Journal, Book and Book series while publications in English language were only considered. The search query is given as TITLE-ABS-KEY (“Tomato” OR “solanum lycopersicum” AND (“metabolomics” OR “proteomics” OR “genomics” OR “Transcriptomics”))) AND PUBYEAR > 2000 AND PUBYEAR < 2024 AND (LIMIT-TO (DOCTYPE, “ar”)) OR LIMIT-TO (DOCTYPE, “re”)) OR LIMIT-TO (DOCTYPE, “ch”)) OR LIMIT-TO (DOCTYPE, “bk”)) OR LIMIT-TO (DOCTYPE, “ed”)) OR LIMIT-TO (DOCTYPE, “dp”))) AND (LIMIT-TO (LANGUAGE, “English”)) (Accessed on 08-05-2024).

This initial search query resulted in a total of 2039 publications. Then the search results were scanned for relevancy of the publication and finally, metadata of 1702 relevant publications was downloaded for bibliometric analysis. The detailed steps involved in the methodology are represented in Fig. 1.

**Fig 1.**
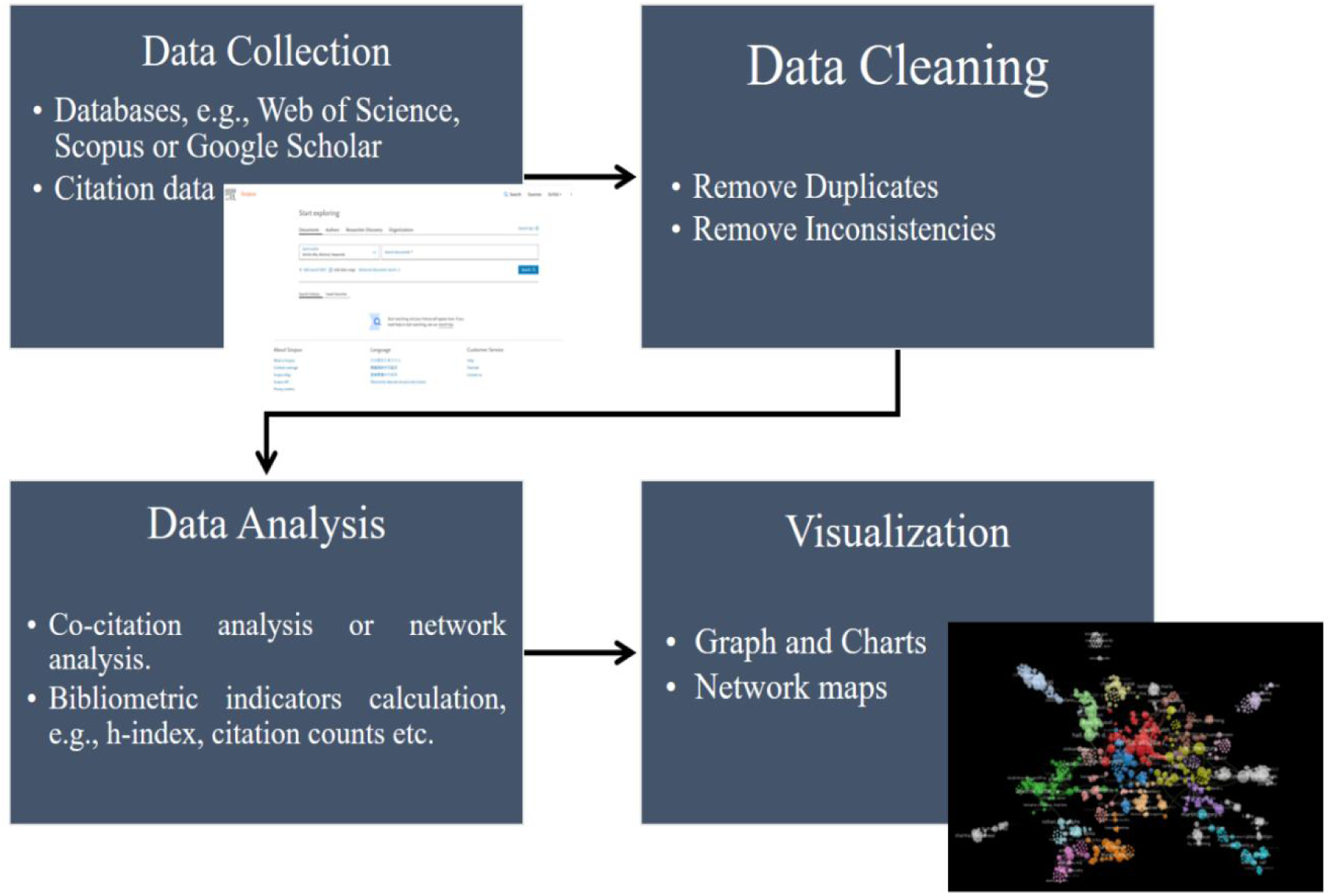
Workflow schematic of bibliometric analysis in scientific literature. The process consists of four major steps: (1) Data Collection through academic databases including Web of Science, Scopus, and Google Scholar for citation data; (2) Data Cleaning to remove duplicates and inconsistencies in the collected data; (3) Data Analysis involving co-citation analysis, network analysis, and calculation of bibliometric indicators; and (4) Visualization of results through graphs, charts, and network maps for effective data representation

### Data analysis

A bibliometric analysis was conducted to identify the shifting landscape of various -omics approaches, i.e., genomics, transcriptomics, proteomics, and metabolomics. This analysis focuses on multiple aspects, including scientific output (e.g. authors and country of origin), research impact (measured by total citations and h-index), prevalent research categories, prominent journals, the evolution of scientific terminology, and collaborative efforts among authors. Data required for this study, including article metadata such as titles, authorship details, abstracts, keywords, subject classifications, journal information, publication years, citation counts, and additional relevant metrics, were taken from SCOPUS for thorough analysis. To assess the academic impact of research endeavours, four key statistics were taken into considerations:

i. Total Publications (TP): It provides the total number of research publications attributed to an individual author, a specific country, a particular publisher, or of a particular journal or research area.
ii. Total Citations (TC): This statistic represents the cumulative count of citations of any particular research publication.
iii. h-index: The h-index is a numerical indicator denoting the number of papers that have received at least the same number of citations as the h-index value.
iv. Average Citation per Publication (ACPP): This index is calculated by dividing the total number of citations by the total number of publications, providing an average citation count per publication for authors, countries, etc.

Additionally, by using the metadata from publications, network visualization using various data attributes was carried out to gain deeper insights. To elucidate the temporal evolution of scientific terminology and the extent of author collaboration within the research domain, network visualizations were conducted utilizing a comprehensive counting algorithm within the VOSviewer software platform (van Eck and Waltman, 2010), as suggested by Biscaro and Giupponi (2014).

i. Keyword Co-occurrence (KC): This network visualization method serves as a tool for mapping scientific networks, wherein each node represents a keyword, and connecting links indicate the co-occurrence of keywords. The size of a node corresponds to its frequency, signifying its significance within the network. Likewise, the thickness of a link reflects the strength of the relationship between paired keywords. This network not only unveils clusters of related knowledge within the field but also elucidates the connections among these clusters. Consequently, it offers enhanced clarity regarding the structure of knowledge, the evolution of trending topics, and the trajectories of the scientific domain (Radhakrishnan et al., 2017; Zhuang et al., 2016).
ii. Co-authorship Analysis (CA): This analytical approach delves into collaborative research endeavors involving two or more researchers from diverse institutions and/or countries. Here, nodes represent individual authors, with their sizes corresponding to the number of articles authored, while links denote co-authorship relationships, with their thickness reflecting the number of co-authored articles. By examining the structure of the scientific community through co-authorship relationships, this method unveils insights into authors’ collaborative patterns and their shared interests within the research field. Previous studies have explored both co-authorship and keyword co-occurrence analyses as network-based approaches (Chen, 2006, 2004; Zhang et al., 2012).

## Result and discussion

### Characteristics of publication outputs

A total of 1702 documents related to genomics, transcriptomics, proteomics and metabolomics in Tomato research were collected (accessed date: Apr 25, 2024). These documents were classified as articles (1439), reviews (164), Book Chapters (77), Editorial (12), Books (7) and Data Paper (3), as displayed in Fig. 2. The yearly publications, citations, and cumulative publications from 2000 to 2023 are presented in chronological order in Fig. 2. The publication rate (articles per year) saw a significant increase after 2017, from 49.88 (2001–2017) to 142.33 (2018–2023), indicating a growing prominence of -omics technologies in tomato research in the recent years.

**Fig 2.**
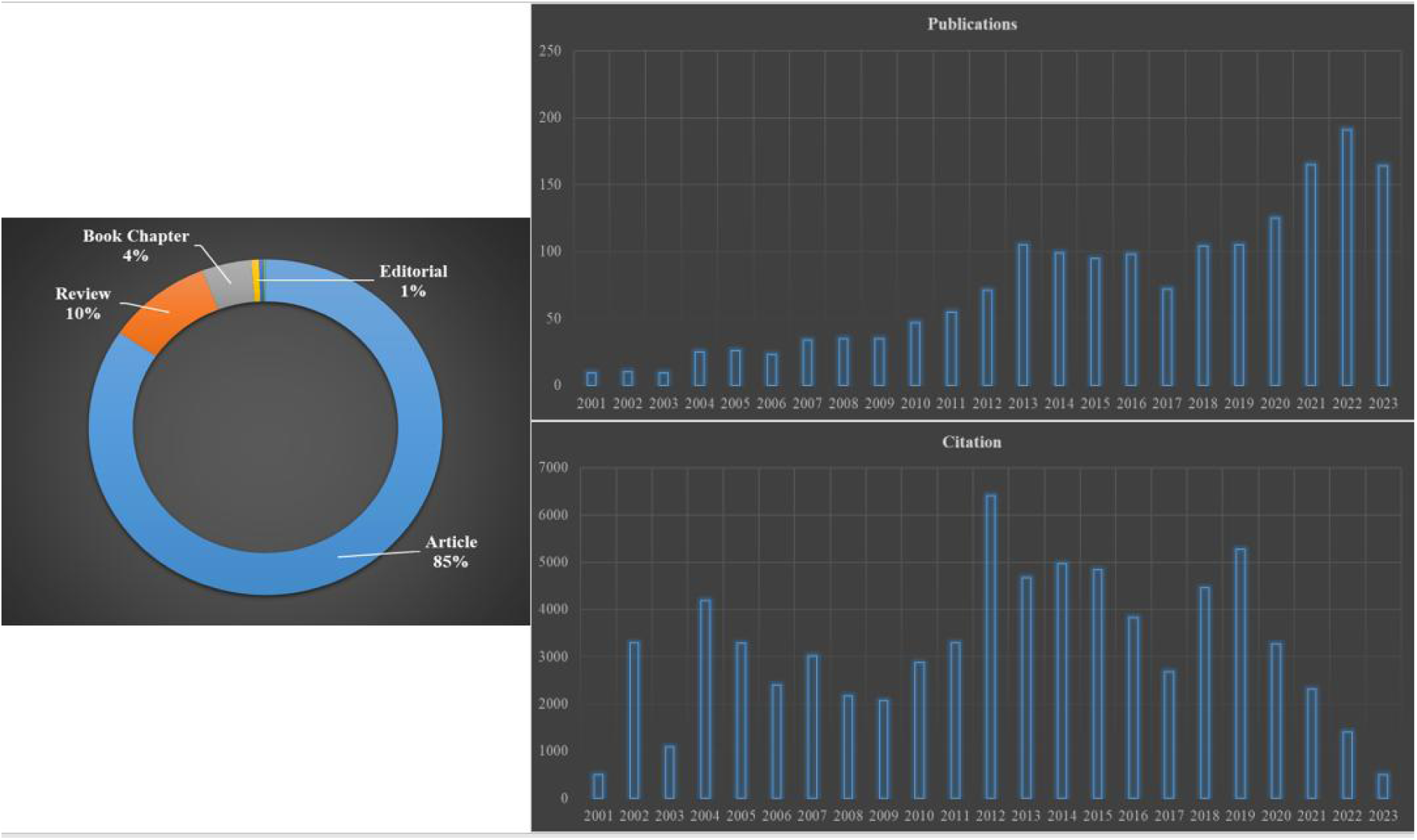
**Document type and publication and citation tend from 2001-2023**

### Publication impact analysis

Publications that are frequently read and referred to are considered to have a significant impact in a given field (Lipsman et al., 2014; Zyoud et al., 2015). Table 1 lists the top 10 most cited publications in tomato omics research. The most cited paper is Kozomara et al. (2019) on miRBase with 2,570 citations and highest FWCI of 97.55, followed by the tomato genome sequencing paper (2012) with 2,418 citations. The topics range from genomics to metabolomics, with publications spanning 2002-2019. Notable is the high impact of methodology papers (miRNA database, genome sequencing) and fundamental research in carotenoid biosynthesis and plant defense. The publications are cited between 2547 (Kozomara et al, 2019) to 444 (Giovannoni et al., 2007) times, indicating their popularity and relevance in the field. The most-cited publication titled ‘‘MiRBase: From microRNA sequences to function’’ by Kozomara et al, 2019 is over microRNA identification in tomato and other crops. The microRNAs are very crucial factor in current research and are reported to modify plant organ morphogenesis, growth, development, hormone secretion, signal transduction, and capacity of plants to react towards external stress and environmental factors (Liu et al., 2009a; Yokotani et al., 2009; Naqvi et al., 2012; Dong et al, 2022). Next most cited paper is over tomato genome sequence (Tomato Genome Consortium, 2012) which provides details of complete genome sequence of tomato bringing a revolution in the field of tomato genomics. All of these publications were published in high-impact factor journals (according to Journal Citation Report: 2022), ranging from 8.7 (Plant Physiology) to 64.5 (Cell) and 64.8 (Nature). Rest of them were also having very high impact factor, i.e. Molecular Plant Pathology (9.3), Current Opinion in Plant Biology (9.5), Plant Journal (11.6), Progress in Lipid Research (13.6), Nucleic Acids Research (14.9), Nature Genetics (30.8) and Nature Biotechnology (46.9). This trend indicates that articles on advanced technology are typically published in high-impact journals and tend to receive more citations compared to those in low-impact journals (Calcagno et al., 2012; Van Noorden, 2017). However, citation counts can differ on platforms like Google Scholar, which includes citations from non-journal sources as well (Yang and Meho, 2006), potentially affecting the rankings.

**Table 1.** Highly cited publications in field of omics research in tomato (*Solanum lycopersicum*) research.

| S. No. | Publication | YoP: Year of Publication | TC; Total Citation | ACPY: Average Citation Per Year | FWCI |
| --- | --- | --- | --- | --- | --- |
| 1. | Kozomara A, Birgaoanu M, Griffiths-Jones S. MiRBase: From microRNA sequences to function. Nucleic Acids Research, 47(D1), pp. D155-D162 | 2019 | 2570 | 428.34 | 97.55 |
| 2. | Sato S, Tabata S, Hirakawa H, et al. The tomato genome sequence provides insights into fleshy fruit evolution. Nature, 485(7400), pp. 635-641<sup>#</sup> | 2012 | 2418 | 186 | 41.54 |
| 3. | Liu Y, Schiff M, Dinesh-Kumar SP. Virus-induced gene silencing in tomato. Plant Journal, 31(6), pp. 777-786 | 2002 | 1240 | 53.91 | 6.41 |
| 4. | Fraser PD, Bramley PM. The biosynthesis and nutritional uses of carotenoids. Progress in Lipid Research, 43(3), pp. 228-265 | 2004 | 1119 | 53.28 | 4.28 |
| 5. | Dixon RA, Achnine L, Kota P, et al. The phenylpropanoid pathway and plant defence - A genomics perspective. Molecular Plant Pathology, 3(5), pp. 371-390. | 2002 | 1074 | 46.69 | 4.49 |
| 6. | Kim S, Park M, Yeom S-I, et al. Genome sequence of the hot pepper provides insights into the evolution of pungency in Capsicum species. Nature Genetics, 46(3), pp. 270-278 | 2014 | 754 | 68.54 | 14.97 |
| 7. | Shimatani Z, Kashojiya S, Takayama M, et al. Targeted base editing in rice and tomato using a CRISPR-Cas9 cytidine deaminase fusion. Nature Biotechnology, 35(5), pp. 441-443 | 2017 | 591 | 73.87 | 23.36 |
| 8. | Zhu G, Wang S, Huang Z, et al. Rewiring of the Fruit Metabolome in Tomato Breeding. Cell, 172(1-2), pp. 249-261.e12. | 2018 | 581 | 83 | 15.28 |
| 9. | Moco S, Bino RJ, Vorst O, et al. A liquid chromatography-mass spectrometry-based metabolome database for tomato. Plant Physiology, 141(4), pp. 1205-1218. | 2006 | 466 | 24.53 | 9.16 |
| 10 | Giovannoni JJ. Fruit ripening mutants yield insights into ripening control. Current Opinion in Plant Biology, 10(3), pp. 283-289. | 2007 | 444 | 24.67 | 3.07 |

**\*FWCI:** Field-Weighted Citation Impact shows how well cited this document is when compared to similar documents. A value greater than 1.00 means the document is more cited than expected according to the average. It takes into account: The year of publication, Document type, and Disciplines associated with its source. The FWCI is the ratio of the document's citations to the average number of citations received by all similar documents over a three-year window. Each discipline makes an equal contribution to the metric, which eliminates differences in researcher citation behavior.
<sup>#</sup> cited as Tomato Genome Consortium, 2012

### Author statistics and co-authorship analysis

A total of 8103 authors have contributed to 1702 publications in this field of research in the 21^st^ century, out of which 159 authors were having 5 or more publications. The average number of authors per publication was found to be approximately 5. Table 2 ranks authors by their contributions to tomato omics research. Fernie, A.R. leads with 39 publications and 2,754 citations (h-index 26), followed by Lucini, L. with 26 publications. Notable is the high citation impact of authors like Giovannoni, J.J. (ACPP 301.25) about tomato genome sequencing followed by Rose J.K.C. (ACPP 274.39). Many top authors’ most cited works are linked to the tomato genome project, indicating its fundamental importance to the field.

**Table 2.** Top contributing authors worked in –omics assisted tomato (*Solanum lycopersicum*) research.

| No. | Author | TP | TC | h-index | ACPP | APP | TC of APP |
| --- | --- | --- | --- | --- | --- | --- | --- |
| 1. | Fernie, A.R. | 39 | 2754 | 26 | 70.62 | DOI: 10.1016/j.cell.2017.12.019 | 581 |
| 2. | Lucini, L. | 26 | 609 | 13 | 23.42 | DOI: 10.3389/fpls.2019.00047 | 91 |
| 3. | Thannhauser, T.W. | 22 | 3273 | 14 | 148.77 | DOI: 10.1038/nature11119 | 2418 |
| 4. | Hall, R.D. | 19 | 1618 | 15 | 85.16 | DOI: 10.1104/pp.105.068130 | 443 |
| 5. | Ezura, H. | 18 | 1702 | 13 | 94.56 | DOI: 10.1038/nbt.3833 | 591 |
| 6. | Rose, J.K.C. | 18 | 4939 | 17 | 274.39 | DOI: 10.1038/nature11119 | 2418 |
| 7. | Fei, Z. | 16 | 3862 | 14 | 241.38 | DOI: 10.1038/nature11119 | 2418 |
| 8. | Giovannoni, J.J. | 16 | 4820 | 13 | 301.25 | DOI : 10.1038/nature11119 | 2418 |
| 9. | Aharoni, A. | 15 | 934 | 12 | 62.27 | DOI: 10.1073/pnas.1912130117 | 301 |
| 10. | Tohge, T. | 15 | 1059 | 12 | 70.60 | DOI: 10.1073/pnas.1309606110 | 279 |

TP: Total Publication; TC: Total Citation; ACPP: Average Citations Per Publication; Authors' Popular Publication (APP); TC of APP

The bibliometric analysis reveals complex patterns of scientific collaboration through a network visualization containing 25 distinct author clusters with 6039 links and a total link strength of 7874 (Fig 3). Each cluster, represented by different colors, indicates groups of researchers who frequently collaborate, with the size of each node reflecting an author’s publication output and connecting lines demonstrating co-authorship relationships. Fernie, Alisdair R. emerges as a central figure in the network, positioned at the core with numerous strong connections radiating outward, suggesting their role as a key collaborator and influential researcher in the field. The largest cluster (Cluster 1) contains 60 authors, indicating a substantial collaborative group, followed by green and blue clusters with 55 and 50 authors, respectively. Subsequent clusters decreased in size, with Cluster 25 containing 5 authors, representing smaller collaborative networks. Notable collaborative groups included those centered around researchers like Lucini, Luigi and Martin, Gregory B., each forming distinct clusters with their frequent co-authors. The network structure showed both tight-knit research communities (indicated by dense interconnections within clusters) and bridge researchers who connect different groups (shown by nodes linking multiple clusters). The spatial distribution of the network demonstrates both the centrality of certain researchers and the peripheral nature of others, with some clusters being more isolated, suggesting specialized research groups working on specific aspects of the field. The visualization effectively captures the hierarchical nature of research collaboration, from major research hubs to smaller, specialized research teams. This co-authorship network has provided valuable insights into the social structure of scientific collaboration in the field, highlighting both established research communities and the extent of cross-group collaboration.

**Fig 3.**
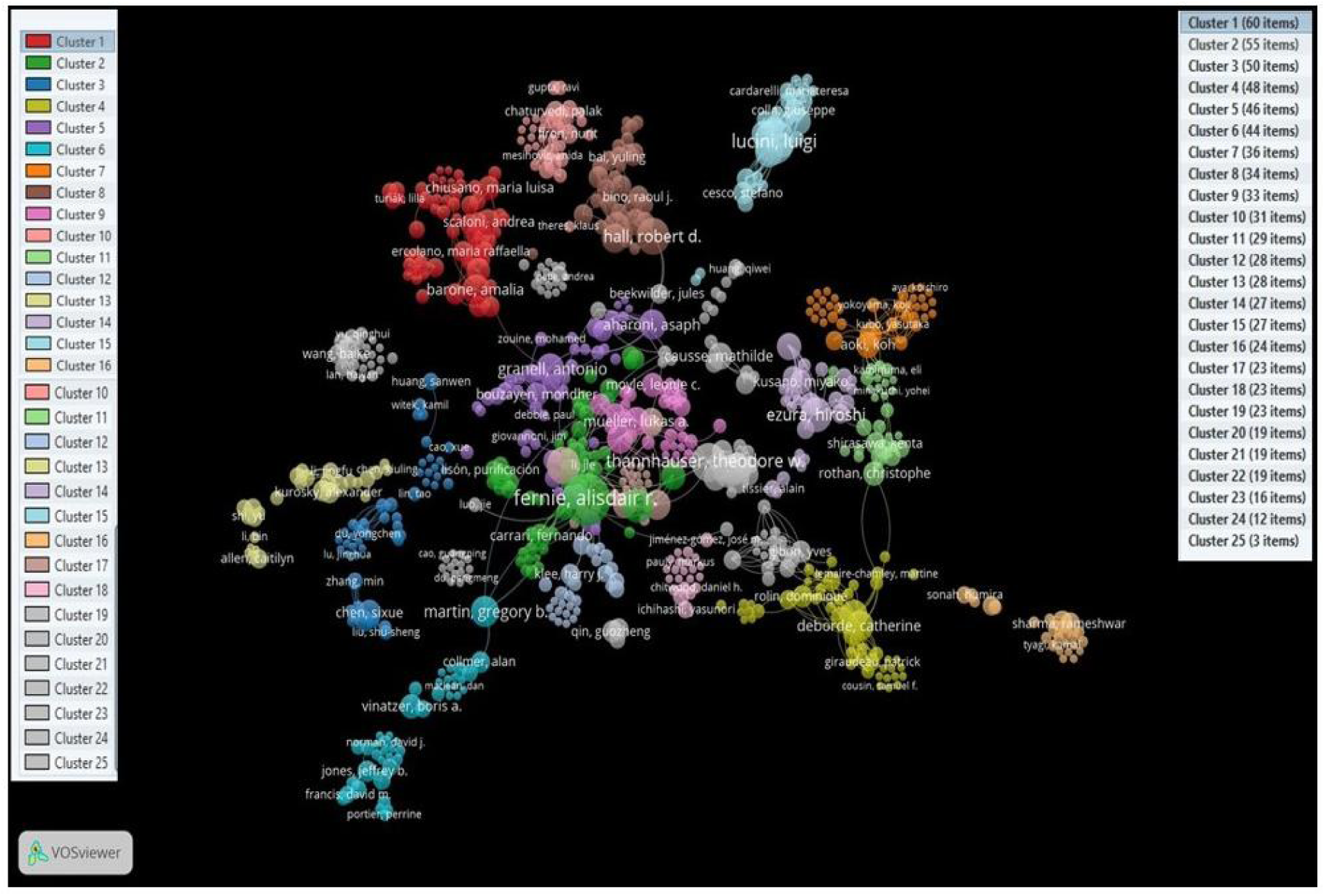
Network visualization of co-authorship patterns in scientific publications generated using VOSviewer. The network comprises 25 distinct clusters (indicated by different colors), with node size representing publication frequency and connecting lines indicating co-authorship strength. Major clusters are centered around key researchers like Fernie, Alisdair R., with cluster sizes ranging from 60 to 5 items as shown in the legend. The spatial distribution and clustering demonstrate collaborative networks and research group formations within the field. The visualization reveals the extent of scientific collaboration and identifies influential research communities through their publication partnerships

### Research hot-spots and keyword co-occurrence analysis

Authors use keywords to indicate the theme and ideas of their study, and analyzing these keywords can help identify and understand the development and research trends in a field over time (Yu et al., 2017). A keywords co-occurrence analysis was conducted through VOSviewer using full counting method (Fig 4). A total of 4087 keywords were identified and keywords with minimum 5 occurrences were further considered, leading to 217 keywords for analysis. The network visualization revealed the interconnected landscape of tomato research, with distinct clusters representing major research domains. The central position of “tomato” and “metabolomics” indicated their pivotal role in current research focus. Key research areas radiate outward, including genomics (red cluster), metabolite profiling (blue cluster), and plant immunity (purple cluster). Disease-related research is prominently represented through pathogens like *Ralstonia solanacearum* and *Pseudomonas syringae*, highlighting the significance of plant protection studies. The network also demonstrated strong connections between stress responses, secondary metabolism, and fruit quality traits, suggesting integrated approaches in tomato improvement research. Modern research tools and approaches such as mass spectrometry, machine learning, and targeted proteomics are well-represented, indicating the field’s technological advancement. The visualization effectively captures the multidisciplinary nature of contemporary tomato research, from fundamental genomics to applied aspects like biocontrol and fruit quality enhancement. Overall, this keyword co-occurrence network visualization offers a comprehensive overview of the major research themes, technologies, and areas of interest within the field of tomato research, as well as their interconnections and relative prominence based on bibliometric data analysis.

**Fig 4.**
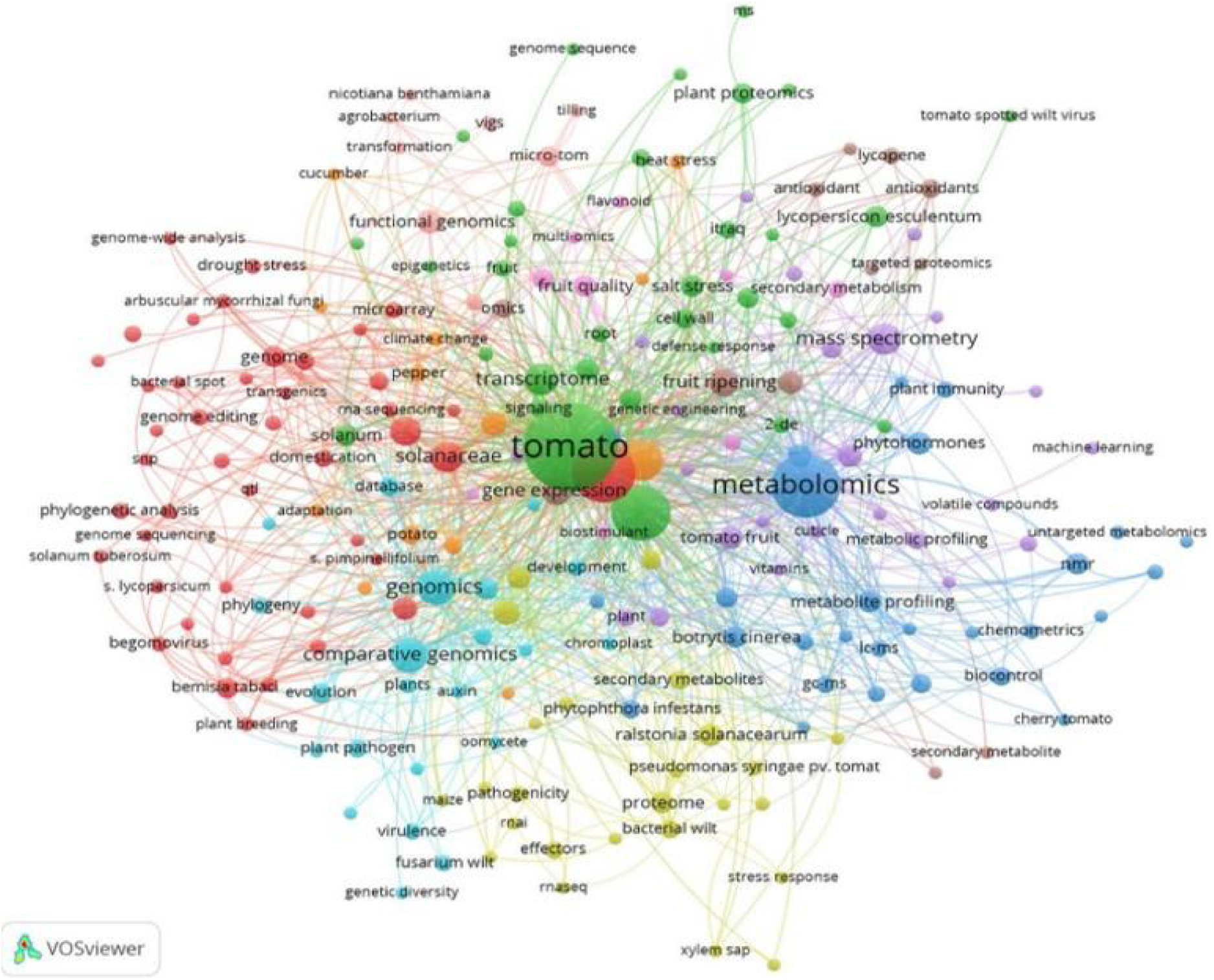
Network visualization of research topic co-occurrence in tomato-related scientific literature. The map was generated using VOSviewer, displaying the interconnected research themes in tomato studies. Node size indicates occurrence frequency, while different colors represent distinct research clusters. Major research areas include metabolomics, genomics, plant immunity, and stress responses. The connecting lines represent the strength of relationships between research topics, with tomato as the central node showing strong connections to various research domains including fruit quality, gene expression, proteomics, and pathogen resistance studies. This network analysis reveals the multidisciplinary nature and key research priorities in tomato research.

### Subject categories’ co-occurrence and key journal ranking analysis

The analysis of research categories (Fig 5a) revealed that ‘Biochemistry, Genetics and Molecular Biology’ dominated the field with the highest number of publications (1054), citations, h-index, and average citations per publication. This was followed by “Agricultural and Biological Sciences” with 877 publications as the second most productive category, demonstrating the strong integration of fundamental and applied research in tomato -omics studies. Medicine, Immunology and Microbiology, and Chemistry show comparatively lower but notable contributions, indicating the broad interdisciplinary nature of tomato research.

**Fig 5.**
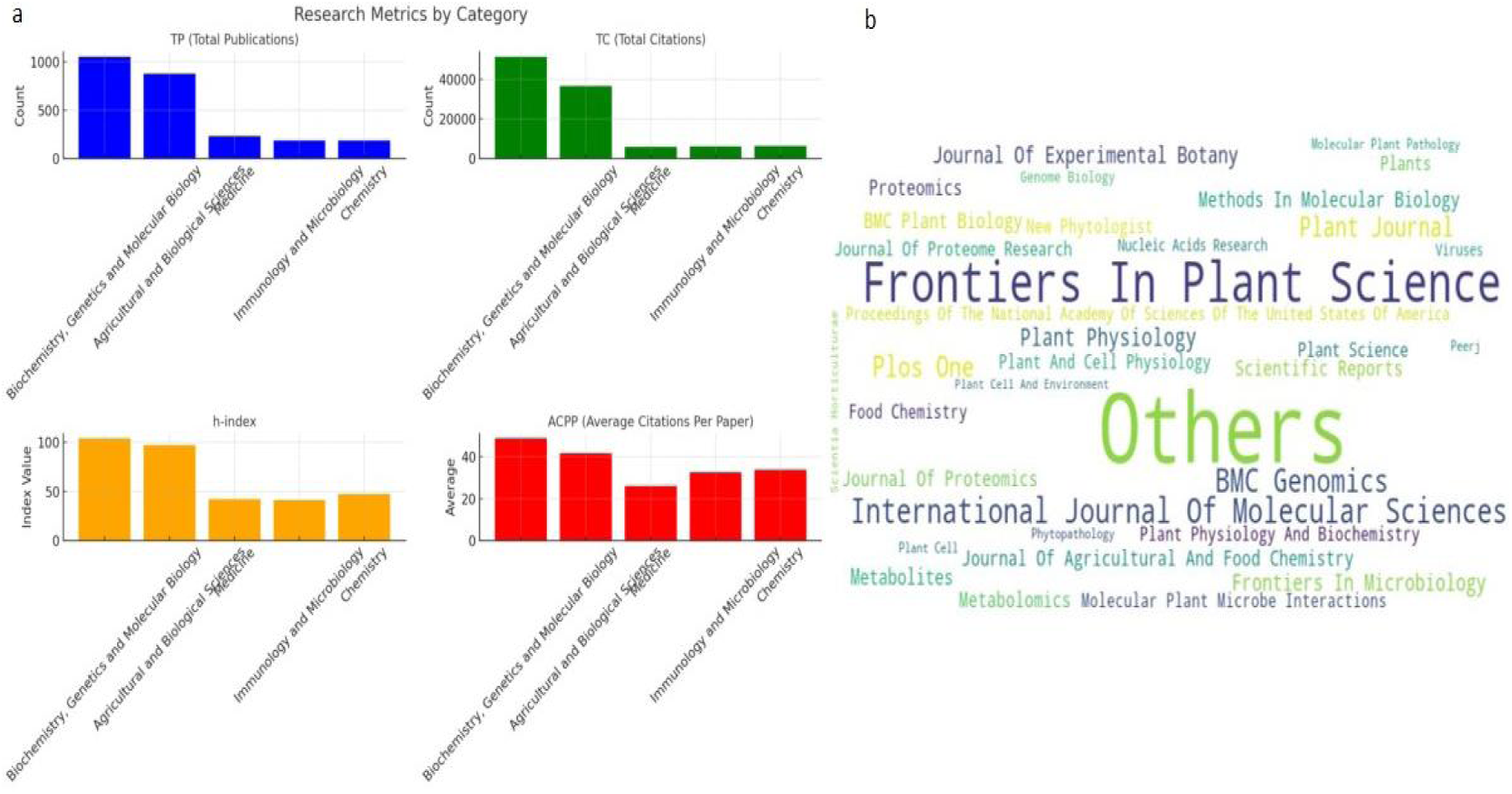
Publication metrics in tomato -omics research (2001-2023), a. Distribution of research output across major scientific categories showing total publications, citations, h-index, and average citations per publication. Biochemistry, Genetics and Molecular Biology emerged as the dominant research category, followed by Agricultural and Biological Sciences. B. Word cloud visualization of journal titles publishing tomato -omics research, with size of text proportional to publication frequency. ‘Frontiers in Plant Science’ and ‘Others’ represent major publication outlets, highlighting the diverse range of journals contributing to tomato -omics literature

The journal distribution analysis (Fig 5b) visualized as a word cloud highlights “Frontiers in Plant Science” as a primary publication venue (92 articles) for tomato -omics research. The prominence of various journals including ‘International Journal of Molecular Sciences (67)’ and ‘BMC Genomics (58)’ followed by “Plant Physiology”. The specialized journals in proteomics, metabolomics, and molecular biology published diverse methodological approaches in the field. The significant presence of “Others” suggested a wide distribution of publications across numerous journals, indicating the broad impact and multidisciplinary nature of tomato -omics research. This bibliometric analysis effectively captured both the dominant research categories and the diverse publication landscape in tomato-omics studies over the past two decades (2001-2023), highlighting its evolution as a multifaceted research area spanning both basic and applied sciences.

Table 3 presents the characteristics of the top ten journals that have published the most research from 2001 to 2023. The statistical results showed that the journal ‘Frontiers in Plant Science (92)’ published the highest number of articles, followed by . ‘Frontiers in Plant Science’ is most cited plant science journal which advances understanding of plant biology for sustainable food security, functional ecosystems and human health. Among the top 10 journals, ‘Plant Journal’ was having highest ‘ACPP’ while ‘Metabolites’ was having lowest (13.19) (Table 3). Further when we look for different ‘Document Type’, then we can observe that they were distributed in three categories, Article, Review and Editorial. Here, only 3 journals published editorial articles.

**Table 3.**
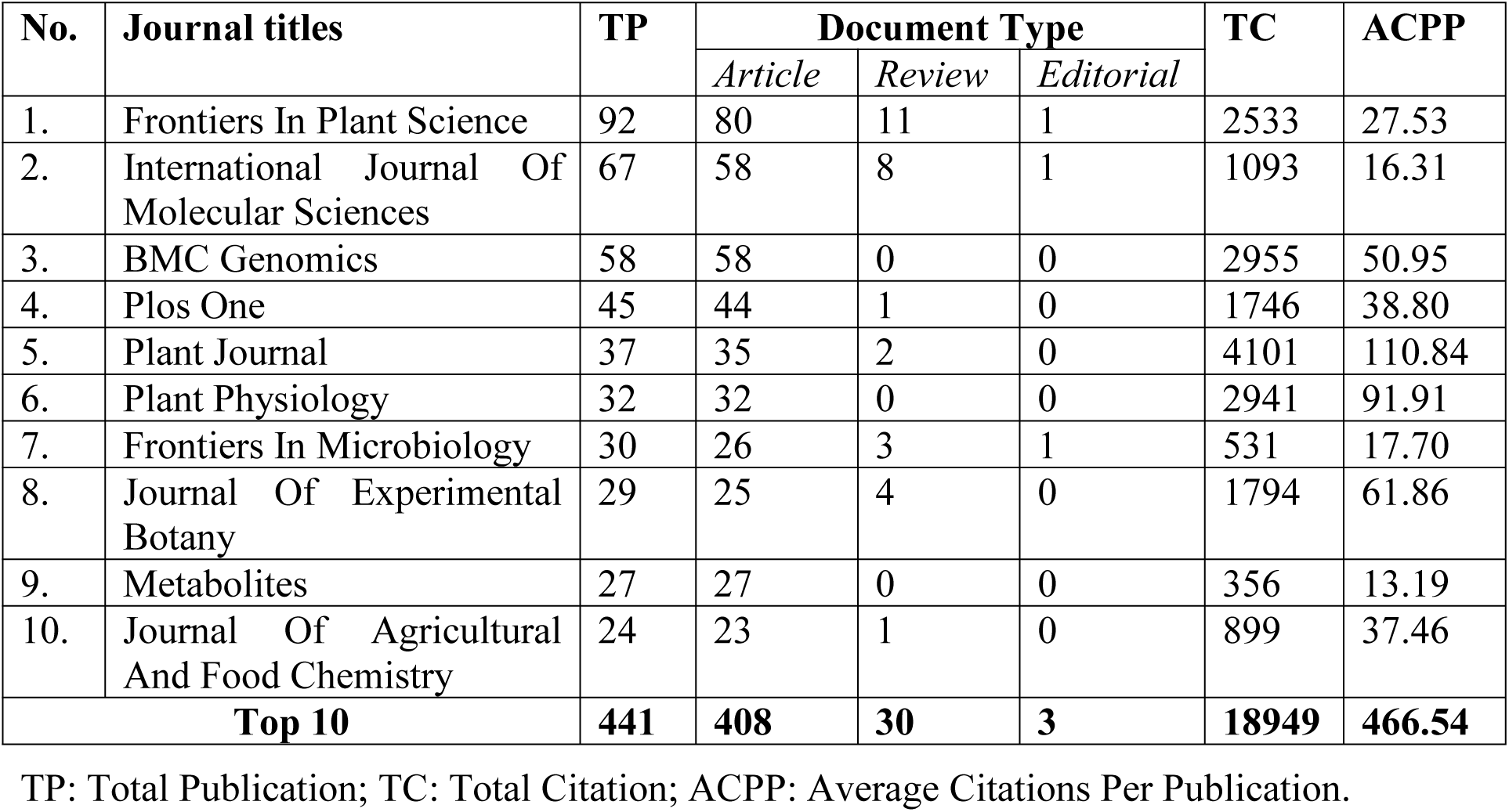
Top 10 journals which publishes studies on -omics research in Tomato (*Solanum lycopersicum*)

| No. | Journal titles | TP | Document Type |  |  | TC | ACPP |
| --- | --- | --- | --- | --- | --- | --- | --- |
|  |  |  | Article | Review | Editorial |  |  |
| 1. | Frontiers In Plant Science | 92 | 80 | 11 | 1 | 2533 | 27.53 |
| 2. | International Journal Of Molecular Sciences | 67 | 58 | 8 | 1 | 1093 | 16.31 |
| 3. | BMC Genomics | 58 | 58 | 0 | 0 | 2955 | 50.95 |
| 4. | Plos One | 45 | 44 | 1 | 0 | 1746 | 38.80 |
| 5. | Plant Journal | 37 | 35 | 2 | 0 | 4101 | 110.84 |
| 6. | Plant Physiology | 32 | 32 | 0 | 0 | 2941 | 91.91 |
| 7. | Frontiers In Microbiology | 30 | 26 | 3 | 1 | 531 | 17.70 |
| 8. | Journal Of Experimental Botany | 29 | 25 | 4 | 0 | 1794 | 61.86 |
| 9. | Metabolites | 27 | 27 | 0 | 0 | 356 | 13.19 |
| 10. | Journal Of Agricultural And Food Chemistry | 24 | 23 | 1 | 0 | 899 | 37.46 |
| **Top 10** |  | **441** | **408** | **30** | **3** | **18949** | **466.54** |

TP: Total Publication; TC: Total Citation; ACPP: Average Citations Per Publication.

### Countries cooperation analysis

The United States leads with 450 publications and 30,080 citations (h-index 86), followed by China (310 publications) and Italy (171 publications) (Table 4). Interestingly, while the UK ranks 10th in publication count (102), it has the highest ACPP (115.81), indicating high impact per publication. The data shows strong contributions from both Western and Asian countries in tomato omics research. Overall, the top 10 countries together account for approximately 69% of the total research publications. (Fig 6A). Based on the total citation and h-index statistics, Netherlands and United Kingdom are also producing popular publications that are highly cited.

**Table 4.** Top ten productive countries in omics research in tomato (*Solanum lycopersicum*)

| S. No. | Country/Territory | Total Publications | TC: Total Citation | h-index | ACPP: Average Citations Per Publication |
| --- | --- | --- | --- | --- | --- |
| 1. | United States | 450 | 30080 | 86 | 66.84 |
| 2. | China | 310 | 11816 | 51 | 38.12 |
| 3. | Italy | 171 | 8275 | 46 | 48.39 |
| 4. | Germany | 150 | 9535 | 50 | 63.57 |
| 5. | Spain | 146 | 7856 | 41 | 53.81 |
| 6. | India | 144 | 5194 | 32 | 36.07 |
| 7. | France | 118 | 7353 | 43 | 62.31 |
| 8. | Netherlands | 116 | 9540 | 44 | 82.24 |
| 9. | Japan | 105 | 7590 | 41 | 72.29 |
| 10 | United Kingdom | 102 | 11813 | 44 | 115.81 |

**Fig 6.**
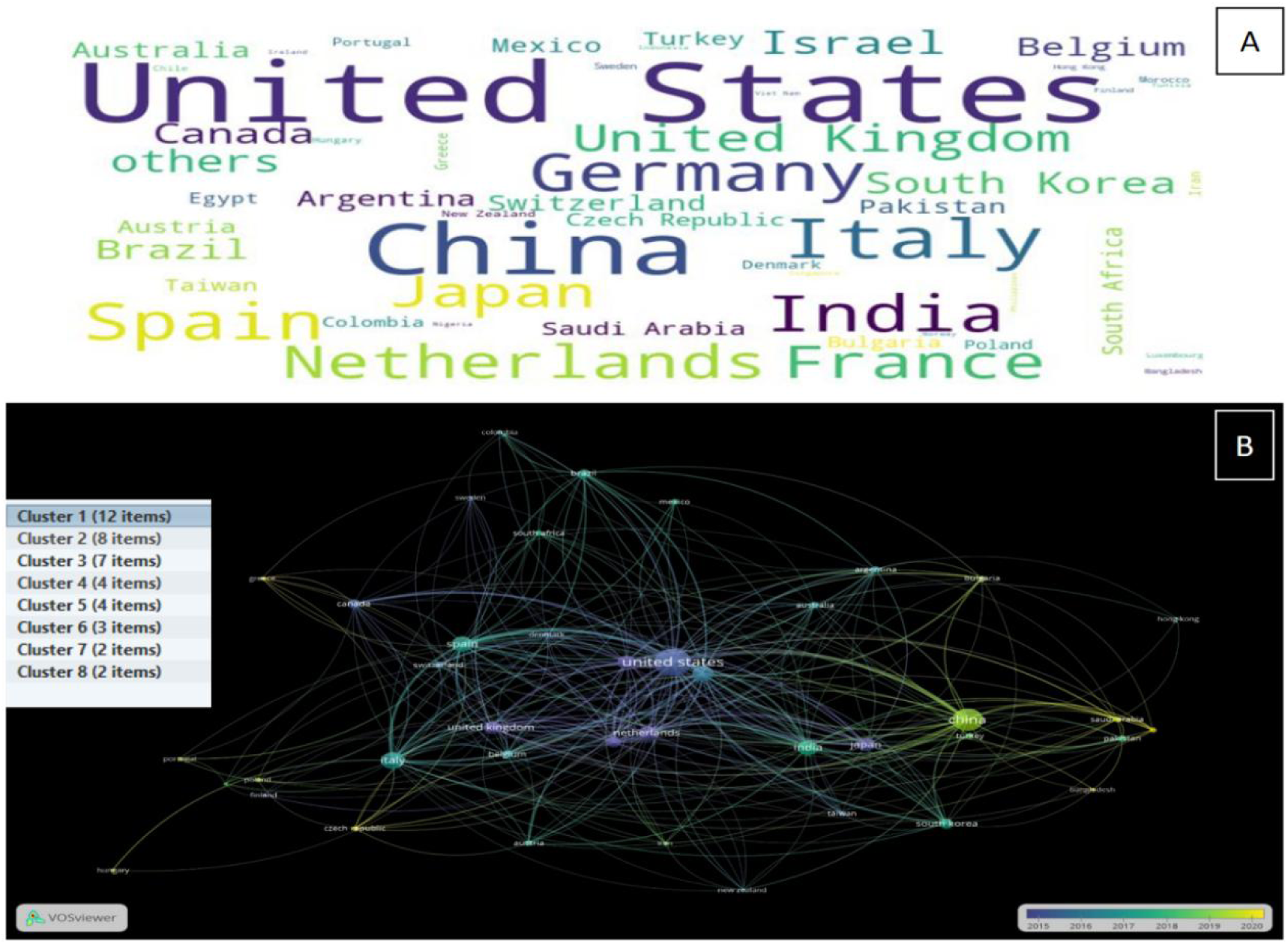
Word cloud depicting countries frequently mentioned in the research collaboration analysis (A) and visualization of the year-wise co-operation network between different countries (B)

Regarding the collaboration in research among the countries, the topmost country i.e. United states showed good cooperation among themselves, with 39 links and a total strength of 393 links (Fig 6.B). A total of 8 clusters were generated where cluster 1 was largest with 12 countries. If we see in years then initially US, UK and other countries were having highest number of publications, though in recent years, paradigm shift is slowly on the way where Asian countries are also having significant no of publications (Fig 6.B).

### Insights from omics research in tomato

The field of tomato research has witnessed significant advancements in genomics, transcriptomics, proteomics, and metabolomics from 2001 to 2023. These cutting-edge technologies have provided valuable insights into the genetic makeup, gene expression patterns, protein profiles, and metabolic processes of tomato plants, contributing to a deeper understanding of their biology and facilitating various applications. The tomato genome was sequenced and published in 2012, marking a significant milestone in tomato research (Tomato Genome Consortium, 2012). This achievement paved the way for comprehensive genetic studies, marker-assisted breeding, and the identification of genes associated with important traits. Several subsequent studies have focused on comparative genomics, exploring the genetic diversity and evolution of tomato species.

Transcriptomic studies have provided insights into the gene expression patterns involved in various aspects of tomato development, fruit ripening, and stress responses. RNA-seq and microarray technologies have been extensively employed to study the transcriptome dynamics during fruit development (Ye et al., 2015), ripening (Zhang et al., 2020), and under abiotic and biotic stress conditions (Amoroso et al, 2023). Proteomic approaches have been utilized to investigate the protein composition and dynamics in tomato tissues and organs. These studies have shed light on the proteome changes associated with fruit development, ripening, and stress responses (Momo et al, 2023; Faurobert et al, 2007; Pontiggia et al, 2019; Wang et al, 2014). Mass spectrometry-based techniques have been instrumental in identifying and quantifying tomato proteins and their post-translational modifications.

Metabolomic studies have focused on profiling and characterizing the diverse metabolites present in tomato fruits and other plant tissues. These analyses have provided valuable information about the metabolic pathways involved in fruit quality, flavor, and nutritional composition (Tang et al, 2020; Zhu et al, 2018; Mun et al, 2021; Ara et al, 2021; Mellidou et al, 2023; Pan et al, 2023). Advanced analytical techniques, such as gas chromatography-mass spectrometry (GC-MS) and liquid chromatography-mass spectrometry (LC-MS), have facilitated the identification and quantification of metabolites in tomato samples. These omics technologies have not only advanced our fundamental understanding of tomato biology but have also contributed to practical applications in breeding programs, crop improvement, and postharvest management. By integrating data from multiple omics platforms, researchers have gained a more comprehensive view of the molecular mechanisms underlying various traits and processes in tomato plants, paving the way for the development of improved tomato varieties with enhanced yield, quality, and resilience.

### Conclusion

Overall, the publication characteristics and trends reflected 1702 publications in omics related research in tomatoes majorly including genomics, genetics, plant physiology, development, fruit quality, stress responses and analytical techniques. The rate of publication increased significantly after 2017, indicating growing implications of omics technologies in recent tomato research. The analysis covered seminal works like the tomato genome sequencing (in 2012) and transcriptomic, proteomic and metabolomic studies that advanced our understanding on tomato biology, development, stress responses and crop improvement applications. The highly cited publications were on topics like microRNA identification, and tomato genome sequencing, published in prestigious journals like Nature, Cell, and Plant Physiology. High citation counts indicated significance and influence of these foundational works in driving subsequent omics research in tomatoes. A total of 8103 authors contributed, with top authors being Fernie AR, Lucini L, Thannhauser TW having the highest number of publications. Co-authorship analysis showed extensive collaborative networks, with clustering patterns indicative of research groups and emerging areas. Keyword analysis highlighted that the biochemistry, genetics, molecular biology and agricultural/biological sciences were the dominant categories of research, though the publications in chemistry, microbiology and immunology had higher citation rates per paper. Co-occurrence patterns showed interconnections between themes like genomics-transcriptomics-proteomics and plant immunity-virulence-effectors. Comprehensive visualization mapped the multi-disciplinary landscape and intersections within tomato omics research. Country collaboration networks revealed research hubs, international collaborations and emerging contributors like Asian nations. Conclusively, the comprehensive bibliometric analysis quantified and visualized the rapid growth, emerging themes, leading contributors, collaboration patterns and multidisciplinary impact of tomato omics research over the past two decades. The findings could guide research priorities, funding, programs and policies related to this economically important crop.

### Perspectives

The bibliometric analysis revealed emerging trends and future directions in tomato omics research. Integration of multiple omics approaches is becoming increasingly important, suggesting a shift toward systems biology perspectives. The rising contribution from Asian countries indicates geographical diversification of research expertise. Emerging research areas have included the application of machine learning to large-scale omics data analysis, stress response mechanisms, and biofortification studies. Future research priorities have to focus on translating omics insights into practical applications for crop improvement, particularly in addressing climate resilience and nutritional enhancement. The strong collaborative networks identified suggest opportunities for expanding international partnerships, especially between established research hubs and emerging contributors. Additionally, the increasing focus on metabolomics and proteomics approaches complements genomic studies, suggesting a more comprehensive understanding of tomato biology is emerging. These trends highlight the need for continued support of interdisciplinary research and technology development in tomato improvement programs.

## Declarations

### Ethics approval and consent to participate

Not applicable

### Availability of data and materials

The datasets during and/or analysed during the current study available from the corresponding author on reasonable request.

### Authors’ contributions

RP and DPS conceptualized the work, collected and analyzed the data and wrote the MS. AK and Renu reviewed and revised the manuscript draft. All authors have read and approved the final manuscript.

## Acknowledgements

Authors are thankful to Indian Council of Agricultural Research, (ICAR), INDIA for funding and Institutional support. DPS is thankful to Network Project on Agricultural Bioinformatics by ICAR, INDIA for funding support in the form of CABin Project.

## References

Amoroso CG, D’Esposito D, Aiese Cigliano R, Ercolano MR. 2023. Comparison of tomato transcriptomic profiles reveals overlapping patterns in abiotic and biotic stress responses. Int J Mol Sci. 24:4061. 10.3390/ijms24044061

Ara T, Sakurai N, Takahashi S, Waki N, Suganuma H, Aizawa K, Matsumura Y, Kawada T, Shibata D. 2021. TOMATOMET: A metabolome database consists of 7118 accurate mass values detected in mature fruits of 25 tomato cultivars. Plant Direct. 5(4):e00318. doi: 10.1002/pld3.318.

Baas J, Schotten M, Plume A, Côté G, Karimi R. 2020. Scopus as a Curated, High-Quality Bibliometric Data Source for Academic Research in Quantitative Science Studies. Quant Sci Stud. 1: 377–386.

Bento-da-Silva EPP, Mendonça SR, de Moraes MG. 2023. Trends and gaps in tomato grafting literature: a systematic approach. Span J Agric Res. 21(3): e0904.

Bhowmik D, Kumar KS, Paswan S, Srivastava S. 2012. Tomato-A Natural Medicine and Its Health Benefits. J Pharmacogn Phytochem. 1:33–43.

Biscaro C, Giupponi C. 2014. Co-authorship and bibliographic coupling network effects on citations. PLoS One. 9, e99502.

Borguini RG, Ferraz Da Silva Torres EA. 2009. Tomatoes and Tomato Products as Dietary Sources of Antioxidants. Food Rev Int. 25:313–325. doi: 10.1080/87559120903155859.

Broadus RN. 1087. Toward a definition of “bibliometrics”. Scientometrics. 12(5-6), 373–379.

Calcagno V, Demoinet E, Gollner K, Guidi L, Ruths D, de Mazancourt C. 2012. Flows of research manuscripts among scientific journals reveal hidden submission patterns. Science. 338, 1065–1069.

Chen C. 2004. Searching for intellectual turning points: Progressive knowledge domain visualization. Proc Natl Acad Sci. 101, 5303–5310.

Chen C. CiteSpace II: 2006. Detecting and visualizing emerging trends and transient patterns in scientific literature. J Am Soc Inf Sci Technol. 57, 359–377.

Collins EJ, Bowyer C, Tsouza A, Chopra M. 2022. Tomatoes: an extensive review of the associated health impacts of tomatoes and factors that can affect their cultivation. Biology (Basel). 11(2):239. doi: 10.3390/biology11020239.

Dong Q, Hu B, Zhang C. 2022. microRNAs and their roles in plant development. Front Plant Sci. 13:824240. doi: 10.3389/fpls.2022.824240.

Faurobert M, Mihr C, Bertin N, Pawlowski T, Negroni L, Sommerer N, Causse M. 2007. Major proteome variations associated with cherry tomato pericarp development and ripening. Plant Physiol. 143:1327–1346. 10.1104/pp.106.092817

Giovannoni JJ. Fruit ripening mutants yield insights into ripening control. 2007. Curr Opin Plant Biol. 10:283–9. doi: 10.1016/j.pbi.2007.04.008.

Harzing AW, Alakangas S. 2016. Google Scholar, Scopus and the Web of Science: a longitudinal and cross-disciplinary comparison. Scientometrics. 106, 787–804.

Kearney PM, Whelton M, Reynolds K, Muntner P, Whelton PK, He J. 2005. Global burden of hypertension: analysis of worldwide data. Lancet. 5:365:217–223. doi: 10.1016/S0140-6736(05)17741-1.

Kozomara A, Birgaoanu M, Griffiths-Jones S. 2019. miRBase: from microRNA sequences to function. Nucleic Acids Res. 47(D1):D155–D162. doi: 10.1093/nar/gky1141.

Kulshrestha Kinjal, Parihar Akarsh. 2020. A decade of tomato transcriptomics: status and perspectives. Int J Curr Microbiol App Sci. 9:2026–2056. doi: 10.20546/ijcmas.2020.903.235

Kumpulainen M, Seppänen M. 2022. Combining Web of Science and Scopus datasets in citation-based literature study. Scientometrics. 127, 5613–5631. 10.1007/s11192-022-04475-7

Li K, Rollins J, Yan E. 2018. Web of Science use in published research and review papers 1997–2017: a selective, dynamic, cross-domain, content-based analysis. Scientometrics. 2018; 115, 1–20.

Lipsman N, Woodside DB, Lozano AM. 2014. Trends in anorexia nervosa research: an analysis of the top 100 most cited works. Eur Eat Disord Rev. 22, 9–14.

Liu B, Li J, Tsykin A, Liu L, Gaur AB. 2009. Exploring complex miRNA-mRNA interactions with Bayesian networks by splitting-averaging strategy. BMC Bioinformatics. 10:408. doi: 10.1186/1471-2105-10-408.

Mellidou I, Koukounaras A, Frusciante S, Rambla JL, Patelou E, Ntoanidou S, Pons C, Kostas S, Nikoloudis K, Granell A, Diretto G, Kanellis AK. 2023. A metabolome and transcriptome survey to tap the dynamics of fruit prolonged shelf-life and improved quality within Greek tomato germplasm. Front Plant Sci. 14:1267340. doi: 10.3389/fpls.2023.1267340

Momo J, Rawoof A, Kumar A, Islam K, Ahmad I, Ramchiary N. 2023. Proteomics of reproductive development, fruit ripening, and stress responses in tomato. J Agric Food Chem. 2023; 71:65–95

Mongeon P, Paul-Hus A. 2016. The journal coverage of Web of Science and Scopus: A comparative analysis. Scientometrics. 106(1), 213–228. 10.1007/s11192-015-1765-5

Mun HI, Kwon MC, Lee N-R, Son SY, Song DH, Lee CH. 2021. Comparing metabolites and functional properties of various tomatoes using mass spectrometry-based metabolomics approach. Front Nutr. 8:659646. doi: 10.3389/fnut.2021.659646

Naik B, Kumar V, Rizwanuddin S, Chauhan M, Choudhary M, Gupta AK, Kumar P, Kumar V, Saris PEJ, Rather MA, Bhuyan S, Neog PR, Mishra S, Rustagi S. 2023. Genomics, proteomics, and metabolomics approaches to improve abiotic stress tolerance in tomato plant. Int J Mol Sci. 24(3):3025. doi: 10.3390/ijms24033025.

Naqvi AR, Sarwat M, Hasan S, Roychodhury N. 2012. Biogenesis, functions and fate of plant microRNAs. J Cell Physiol. 227, 3163–3168. doi: 10.1002/jcp.24052

Pan F, Zhang Q, Zhu H, Li J, Wen Q. 2023. Transcriptome and metabolome provide insights into fruit ripening of cherry tomato (*Solanum lycopersicum* var. cerasiforme). Plants. 12:3505. 10.3390/plants12193505

Pontiggia D, Spinelli F, Fabbri C et al. 2019. Changes in the microsomal proteome of tomato fruit during ripening. Sci Rep. 9, 14350. 10.1038/s41598-019-50575-5

Pranckutė R. 2021. Web of Science (WoS) and Scopus: The titans of bibliographic information in today’s academic world. Publications. 9(1):12. 10.3390/publications9010012

Pritchard A. 1969. Statistical bibliography or bibliometrics?. J Doc. 25(4), 348–349.

Radhakrishnan S, Erbis S, Isaacs JA, Kamarthi S. 2017. Novel keyword co-occurrence network-based methods to foster systematic reviews of scientific literature. PLoS One. 12, e0172778.

Rao AV, Agarwal S. 1999. Role of lycopene as antioxidant carotenoid in the prevention of chronic diseases: a review. Nutr Res. 19:305–323. doi: 10.1016/S0271-5317(98)00193-6.

Singh DP, Bisen MS, Prabha R, et al. 2023. Untargeted Metabolomics of Alternaria solani-Challenged Wild Tomato Species *Solanum cheesmaniae* Revealed Key Metabolite Biomarkers and Insight into Altered Metabolic Pathways. Metabolites. 13(5):585. doi: 10.3390/metabo13050585.

Singh DP, Maurya S, Yerasu SR et al. 2023. Metabolomics of early blight (*Alternaria solani*) susceptible tomato (*Solanum lycopersicum*) unfolds key biomarker metabolites and involved metabolic pathways. Sci Rep. 13, 21023. 10.1038/s41598-023-48269-0

Singh DP, Rai N, Farag MA, Maurya S, Yerasu SR, Bisen MS, Prabha R, Shukla R, Behera TK. 2024. Metabolic diversity, biosynthetic pathways, and metabolite biomarkers analysed via untargeted metabolomics and the antioxidant potential reveal for high temperature tolerance in tomato hybrid. Plant Stress. 11 : 100420.

Tang H, Zhang X, Gong B, Yan Y, Shi Q. 2020. Proteomics and metabolomics analysis of tomato fruit at different maturity stages and under salt treatment. Food Chem. 311, 126009.

Tilesi F, Lombardi A, Mazzucato A. 2021. Scientometric and methodological analysis of the recent literature on the health-related effects of tomato and tomato products. Foods. 10(8):1905. 10.3390/foods10081905

Tomato Genome Consortium. 2012. The tomato genome sequence provides insights into fleshy fruit evolution. Nature. 485(7400):635–41. doi: 10.1038/nature11119.

Toor RK, Lister CE, Savage GP. 2005. Antioxidant Activities of New Zealand-Grown Tomatoes. Int J Food Sci Nutr. 56:597–605. doi: 10.1080/09637480500490400.

van Eck NJ, Waltman L. 2010. Software survey: VOSviewer, a computer program for bibliometric mapping. Scientometrics. 84, 523–538. 10.1007/s11192-009-0146-3.

Van Noorden R. 2017. The science that’s never been cited. Nature. 552, 162–165.

Viuda-Martos M, Sanchez-Zapata E, Sayas-Barberá E, Sendra E, Pérez-Álvarez JA, Fernández-López J. 2014. Tomato and tomato byproducts. human health benefits of lycopene and its application to meat products: a review. Crit Rev Food Sci Nutr. 54:1032–1049. doi: 10.1080/10408398.2011.623799.

Wang Y, Wang W, Cai J et al. 2014. Tomato nuclear proteome reveals the involvement of specific E2 ubiquitin-conjugating enzymes in fruit ripening. Genome Biol. 15, 548.

Wouters P, Thelwall M, Kousha K, Waltman L, de Rijcke S, Rushforth A, Franssen T. 2015. The metric tide: literature review (supplementary report I to the independent review of the role of metrics in research assessment and management); HEFCE: Bristol, UK, 2015.

Yang K, Meho LI2006. Citation analysis: a comparison of Google Scholar, Scopus, and Web of Science. Proc Am Soc Inf Sci Technol. 43, 1–15.

Ye J, Hu T, Yang C, Li H, Yang M, Ijaz R, Ye Z, Zhang Y. 2015. Transcriptome profiling of tomato fruit development reveals transcription factors associated with ascorbic acid, carotenoid and flavonoid biosynthesis. PLoS One. 10(7):e0130885. doi: 10.1371/journal.pone.0130885.

Yokotani N, Nakano R, Imanishi S, Nagata M, Inaba A. 2009. Ripening-associated ethylene biosynthesis in tomato fruit is autocatalytically and developmentally regulated. J Exp Bot. 60, 3433–3442. doi: 10.1093/jxb/erp185.

Yu D, Xu Z, Pedrycz W, Wang W. Information sciences 2017. 1968–2016: A retrospective analysis with text mining and bibliometric. Inform Sci. 418–419, 619–634. 10.1016/j.ins.2017.08.031.

Zhang C, Duan W, Chen K, Zhang B. 2020. Transcriptome and methylome analysis reveals effects of ripening on and off the vine on flavor quality of tomato fruit. Postharvest Biol Technol. 162, 111096.

Zhang J, Xie J, Hou W, Tu X, Xu J, Song F, Wang Z, Lu Z. 2012. Mapping the knowledge structure of research on patient adherence: knowledge domain visualization based co-word analysis and social network analysis. PLoS One. 7, e34497.

Zhu G, Wang S, Huang Z, Zhang S, Liao Q, Zhang C, Lin T, Qin M, Peng M, Yang C, Cao X, Han X, Wang X, van der Knaap E, Zhang Z, Cui X, Klee H, Fernie AR, Luo J, Huang S. 2018. Rewiring of the fruit metabolome in tomato breeding. Cell. 172(1-2):249–261.e12. doi: 10.1016/j.cell.2017.12.019.

Zhu J, Liu W. 2020. A Tale of Two Databases: The Use of Web of Science and Scopus in Academic Papers. Scientometrics. 123, 321–335.

Zhuang Y, Zhang L, Du Y, Chen G. 2016. Current patterns and future perspectives of best management practices research: A bibliometric analysis. J Soil Water Conserv. 71, 98A–104A.

Zyoud SH, Al-Jabi SW, Sweileh WM, Al-Khalil S, Alqub M, Awang R. 2015. Global methaemoglobinaemia research output (1940–2013): a bibliometric analysis. Springerplus. 4, 1–7.

