## Supplementary Table 1 for "Tomato Omics Research: Mapping Bibliometric Footprints of Global Research Trends, Collaboration Networks, and Impact Trajectories"

| | Genomics | | | | Transcriptomics |
|---|---|---|---|---|---|
| Year | Cummulative frequency | % Coverage | | Year | Cummulative frequency |
| 2001 | 3 | 0.48 | | 2004 | 2 |
| 2002 | 7 | 1.03 | | 2005 | 4 |
| 2003 | 12 | 1.67 | | 2006 | 6 |
| 2004 | 17 | 2.40 | | 2007 | 9 |
| 2005 | 23 | 3.23 | | 2008 | 13 |
| 2006 | 30 | 4.19 | | 2009 | 18 |
| 2007 | 38 | 5.27 | | 2010 | 24 |
| 2008 | 47 | 6.51 | | 2011 | 32 |
| 2009 | 57 | 7.91 | | 2012 | 41 |
| 2010 | 68 | 9.49 | | 2013 | 52 |
| 2011 | 81 | 11.25 | | 2014 | 66 |
| 2012 | 95 | 13.23 | | 2015 | 84 |
| 2013 | 111 | 15.42 | | 2016 | 105 |
| 2014 | 129 | 17.83 | | 2017 | 131 |
| 2015 | 148 | 20.47 | | 2018 | 163 |
| 2016 | 168 | 23.34 | | 2019 | 202 |
| 2017 | 190 | 26.43 | | 2020 | 248 |
| 2018 | 214 | 29.73 | | 2021 | 304 |
| 2019 | 239 | 33.21 | | 2022 | 371 |
| 2020 | 266 | 36.86 | | 2023 | 450 |
| 2021 | 293 | 40.63 | | 2024 | 541 |
| 2022 | 321 | 44.49 | | 2025 | 647 |
| 2023 | 349 | 48.40 | | 2026 | 766 |
| 2024 | 377 | 52.30 | | 2027 | 899 |
| 2025 | 405 | 56.17 | | 2028 | 1044 |
| 2026 | 432 | 59.94 | | 2029 | 1199 |
| 2027 | 458 | 63.59 | | 2030 | 1362 |
| 2028 | 483 | 67.09 | | 2031 | 1528 |
| 2029 | 507 | 70.39 | | 2032 | 1694 |
| 2030 | 530 | 73.49 | | 2033 | 1855 |
| 2031 | 550 | 76.37 | | 2034 | 2009 |
| 2032 | 570 | 79.03 | | 2035 | 2153 |
| 2033 | 587 | 81.45 | | 2036 | 2283 |
| 2034 | 603 | 83.65 | | 2037 | 2401 |
| 2035 | 617 | 85.63 | | 2038 | 2504 |
| 2036 | 630 | 87.41 | | 2039 | 2593 |
| 2037 | 641 | 89.00 | | 2040 | 2670 |
| 2038 | 652 | 90.40 | | 2041 | 2735 |
| 2039 | 660 | 91.65 | | 2042 | 2790 |
| 2040 | 668 | 92.74 | | 2043 | 2835 |
| 2041 | 675 | 93.70 | | 2044 | 2873 |
| 2042 | 681 | 94.54 | | 2045 | 2903 |
| 2043 | 687 | 95.28 | | 2046 | 2929 |
| 2044 | 691 | 95.92 | | 2047 | 2949 |
| 2045 | 695 | 96.47 | | 2048 | 2966 |

| | | | | |
|---|---|---|---|---|
| 2046 | 699 | 96.96 | 2049 | 2980 |
| 2047 | 702 | 97.37 | | |
| 2048 | 704 | 97.74 | | |
| 2049 | 707 | 98.05 | | |
| 2050 | 709 | 98.32 | | |

| % Coverage | Year | Proteomics<br>Cummulative frequency | % Coverage | Year |
| --- | --- | --- | --- | --- |
| 0.05 | 2002 | 3 | 0.73 | 2002 |
| 0.12 | 2003 | 7 | 1.63 | 2003 |
| 0.20 | 2004 | 12 | 2.72 | 2004 |
| 0.31 | 2005 | 18 | 4.05 | 2005 |
| 0.44 | 2006 | 26 | 5.67 | 2006 |
| 0.60 | 2007 | 34 | 7.60 | 2007 |
| 0.79 | 2008 | 45 | 9.91 | 2008 |
| 1.04 | 2009 | 57 | 12.63 | 2009 |
| 1.34 | 2010 | 71 | 15.81 | 2010 |
| 1.72 | 2011 | 88 | 19.46 | 2011 |
| 2.18 | 2012 | 106 | 23.59 | 2012 |
| 2.75 | 2013 | 127 | 28.18 | 2013 |
| 3.46 | 2014 | 149 | 33.20 | 2014 |
| 4.32 | 2015 | 174 | 38.55 | 2015 |
| 5.36 | 2016 | 199 | 44.13 | 2016 |
| 6.64 | 2017 | 224 | 49.82 | 2017 |
| 8.18 | 2018 | 250 | 55.48 | 2018 |
| 10.02 | 2019 | 274 | 60.96 | 2019 |
| 12.22 | 2020 | 298 | 66.17 | 2020 |
| 14.81 | 2021 | 320 | 70.99 | 2021 |
| 17.83 | 2022 | 339 | 75.37 | 2022 |
| 21.30 | 2023 | 357 | 79.28 | 2023 |
| 25.23 | 2024 | 372 | 82.70 | 2024 |
| 29.60 | 2025 | 386 | 85.65 | 2025 |
| 34.38 | 2026 | 397 | 88.17 | 2026 |
| 39.50 | 2027 | 407 | 90.30 | 2027 |
| 44.85 | 2028 | 415 | 92.08 | 2028 |
| 50.32 | 2029 | 421 | 93.55 | 2029 |
| 55.77 | 2030 | 427 | 94.76 | 2030 |
| 61.10 | 2031 | 431 | 95.76 | 2031 |
| 66.16 | 2032 | 435 | 96.57 | 2032 |
| 70.89 | 2033 | 438 | 97.24 | 2033 |
| 75.19 | 2034 | 440 | 97.77 | 2034 |
| 79.05 | 2035 | 442 | 98.21 | 2035 |
| 82.45 | 2036 | 444 | 98.56 | 2036 |
| 85.40 | 2037 | 445 | 98.84 | 2037 |
| 87.92 | 2038 | 446 | 99.07 | 2038 |
| 90.06 | 2039 | 447 | 99.25 | 2039 |
| 91.86 | 2040 | 448 | 99.52 | 2040 |
| 93.35 | 2041 | 449 | 99.69 | 2041 |
| 94.59 | 2042 | 450 | 99.87 | 2042 |
| 95.61 | | | | 2043 |
| 96.44 | | | | 2044 |
| 97.12 | | | | 2045 |
| 97.67 | | | | 2046 |

| | |
|---|---|
| 98.12 | 2047 |
| | 2048 |
| | 2049 |
| | 2050 |
| | 2051 |

| Metabolomics Cummulative frequency | % Coverage |
|---|---|
| 4 | 1.13 |
| 7 | 1.81 |
| 10 | 2.57 |
| 14 | 3.44 |
| 17 | 4.42 |
| 22 | 5.51 |
| 27 | 6.74 |
| 32 | 8.11 |
| 38 | 9.64 |
| 45 | 11.33 |
| 52 | 13.19 |
| 60 | 15.24 |
| 69 | 17.47 |
| 78 | 19.89 |
| 89 | 22.50 |
| 100 | 25.29 |
| 111 | 28.26 |
| 124 | 31.39 |
| 136 | 34.66 |
| 150 | 38.05 |
| 164 | 41.54 |
| 177 | 45.08 |
| 177 | 45.08 |
| 192 | 48.65 |
| 206 | 52.22 |
| 219 | 55.75 |
| 233 | 59.21 |
| 246 | 62.57 |
| 259 | 65.80 |
| 271 | 68.89 |
| 283 | 71.81 |
| 294 | 74.55 |
| 304 | 77.10 |
| 313 | 79.47 |
| 321 | 81.65 |
| 329 | 83.64 |
| 336 | 85.45 |
| 343 | 87.09 |
| 349 | 88.57 |
| 354 | 89.90 |
| 359 | 91.09 |
| 363 | 92.15 |
| 367 | 93.10 |
| 370 | 93.93 |
| 373 | 94.68 |

| | |
|---|---|
| 375 | 95.33 |
| 378 | 95.91 |
| 380 | 96.42 |
| 381 | 96.87 |
| 383 | 97.26 |
